# Pick Your Poison: Molecular Evolution of Venom Proteins in Asilidae (Insecta: Diptera)

**DOI:** 10.1101/2020.11.02.365569

**Authors:** Chris M. Cohen, T. Jeffrey Cole, Michael S. Brewer

## Abstract

Robber flies are an understudied family of venomous, predatory Diptera. With the recent characterization of venom from three asilid species, it is possible for the first time to study the molecular evolution of venom genes in this unique lineage. To accomplish this, a novel whole-body transcriptome of *Eudioctria media* was combined with 10 other publicly available asiloid thoracic or salivary gland transcriptomes to identify putative venom gene families and assess evidence of pervasive positive selection. A total of 348 gene families of sufficient size were analyzed, and 33 of these were predicted to contain venom genes. We recovered 151 families containing homologs to previously described venoms, and 40 of these were uniquely gained in Asilidae. Our gene family clustering suggests that many asilidin venom gene families are not natural groupings as originally delimited. Additionally, robber-fly venoms have relatively few sites under positive selection, consistent with the hypothesis that the venom of older lineages are dominated by negative selection acting to maintain toxic function.

## Introduction

Venoms are typically a composition of various neurotoxins, enzymes, ions, and small organic molecules [1]; [2]; [3]; [4]. Venom proteins generally originate via gene duplication of nontoxic proteins that are then selectively expressed in a venom gland (recruitment), and undergo subsequent neofunctionalization [5]; [3]. However, alternative processes like single gene co-option or *de novo* protein evolution are also believed to be significant drivers of venom evolution in some taxa [6]. Venom proteins are often recruited from secretory proteins involved in rapid physiological processes and from those that have stable tertiary structures due to multiple disulfide bonds [7]. Among these polypeptide components, many neurotoxins have abundant inhibitory cysteine knots (ICKs), but venom enzymes typically lack this motif [7].

Venom has independently evolved in at least four lineages of true flies (Insecta: Diptera) [8]. The family Asilidae (AKA robber flies or assassin flies) is unique among these in that the adults, rather than the larvae, are venomous predators. Their closest relatives, Apioceridae and Mydidae, as adults either feed on nectar and other liquids or do not feed at all [9]; [10]. Adult assassin flies have a venom delivery apparatus rather unlike that found in most other venomous arthropods: the proboscis consists of the labium, which forms a tube through which the hypopharynx slides, and elements of the labrum and maxillae, which support the action of the hypopharynx [11]; [12]; venom is produced in a pair of thoracic glands (also called salivary glands) connected via a fused duct to the hypopharynx, and it is this structure that pierces prey and injects the saliva [13]; [12]; [14]; [15].

Robber flies are capable of incapacitating large and dangerous prey quite rapidly, depending on the site of venom injection [16]; [17]; [11]; [12], and they can deliver painful bites to humans [18]; [19]; [20]; [21]. Recent studies have described over 300 venom proteins and have organized novel venoms into fifteen ‘Asilidin’ protein families, using the taxa *Eutolmus rufibarbis, Machimus arthriticus, Dolopus genitalis*, and *Dasypogon diadema* [15]; [22]; [23]. Some of these venoms are found in a single taxon, consistent with observations of variable venom toxicity between species demonstrated by previous authors [24]; [25]; [26]; [27]; [28]; [29]. Toxicity assays of crude venom or isolated components have shown paralytic and other neurotoxic effects in insect and mammalian subjects [24]; [15]; [22]; [30]. Multiple studies also indicate that disulfide-rich peptide scaffolds (e.g., ICKs), such as those found in spiders and scorpions, have been convergently recruited into the robber-fly venom arsenal [15]; [30].

Sunagar & Moran (2015) developed a “two-speed” model of venom evolution in which the venom genes of young lineages (<60 myo) are often under strong positive selection as niche space is explored, while in older lineages (>380 myo) venom genes are typically dominated by purifying selection in order to preserve toxic function [31]. With the recent description of putative venom genes in Asilidae by Drukewitz et al. and Walker et al. (hereafter referred to as DEA and WEA, respectively [15]; [22]), it is now possible to examine the molecular evolution of venom in these flies, particularly the “two-speed” model of Sunagar & Moran. The age of the family Asilidae is estimated to be between 133-158 myo [32]; [33], similar to toxicoferan lizards at 166 myo [34]. Similar to the lizards analyzed in Sunagar and Moran, we expect robber flies to show little evidence of positive selection in their venom genes. To test this hypothesis, we combine a novel whole-body transcriptome of *Eudioctria media* with 10 publicly available asiloid transcriptomes and provide them as input for the recently developed programs FUSTr [35] and toxify [36] to identify putative venom gene families and determine the extent of pervasive positive selection.

## Results

### Quality, Assembly, and Completeness of Transcriptome

Illumina RNA sequencing generated 67.7 million raw reads for the *Eudioctria media* whole-body transcriptome. After preprocessing with trimgalore (v0.3.7), 66.7 million reads (98%) remained. The *Eudioctria media* transcriptome was assembled into 103,352 contigs. BUSCO (v3.0.2) reported that the *E. media* transcriptome had 86% complete single-copy BUSCOs, 5.1% fragmented BUSCOs, and 8.2% missing BUSCOs. The statistics for this and the other transcriptomes are summarized in Table 1.

**Table 1:**
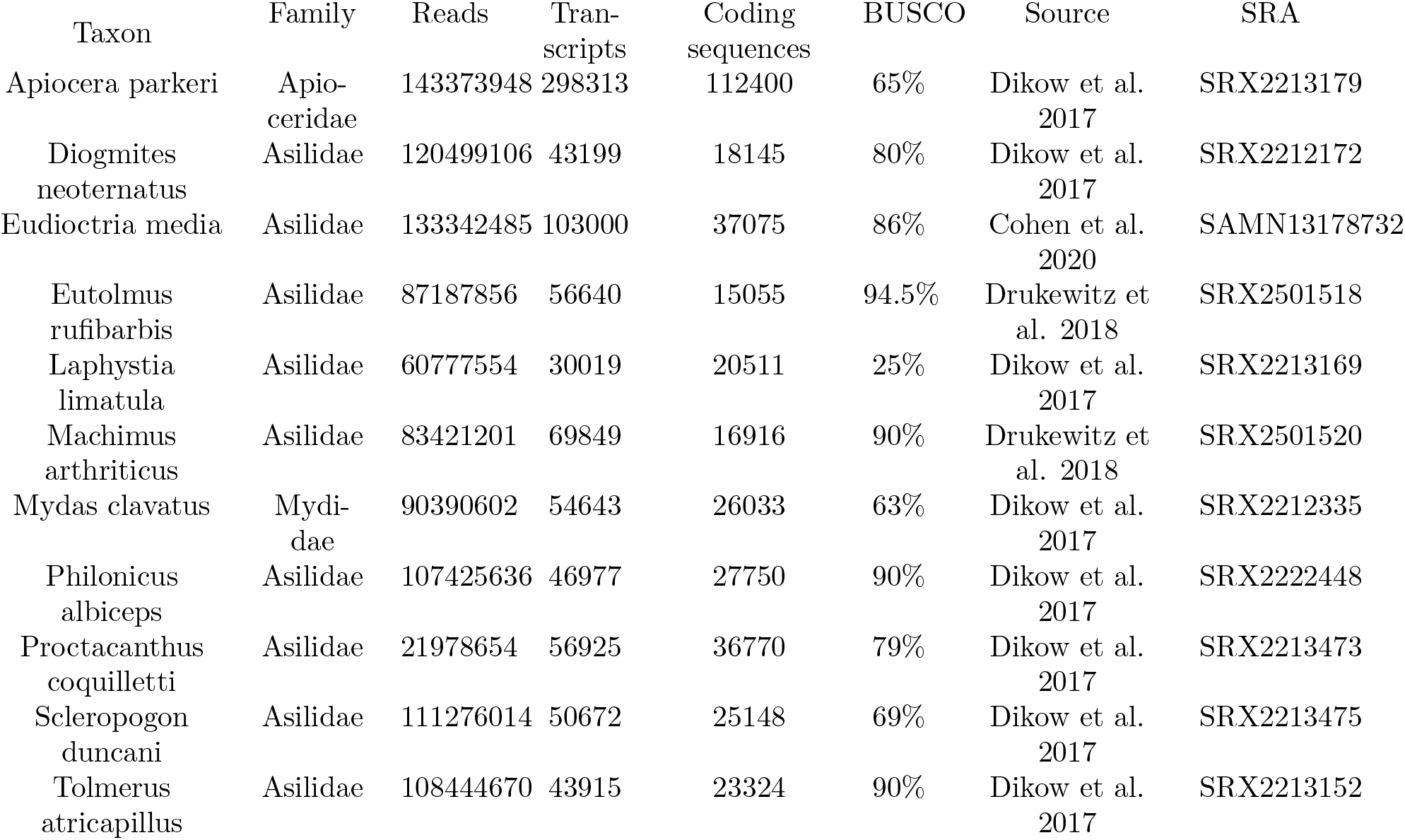
Processed reads, assembled transcripts, BUSCO complete percentage, and source of transcriptome for each taxon included in this study. Values for each transcriptome are derived from their respective publication.

| Taxon | Family | Reads | Transcripts | Coding sequences | BUSCO | Source | SRA |
| --- | --- | --- | --- | --- | --- | --- | --- |
| *Apiocera parkeri* | Apio-<br>ceridae | 143373948 | 298313 | 112400 | 65% | Dikow et al.<br>2017 | SRX2213179 |
| *Diogmites*<br>*neoternatus* | Asilidae | 120499106 | 43199 | 18145 | 80% | Dikow et al.<br>2017 | SRX2212172 |
| *Eudioctria media* | Asilidae | 133342485 | 103000 | 37075 | 86% | Cohen et al.<br>2020 | SAMN13178732 |
| *Eutolmus*<br>*rufibarbis* | Asilidae | 87187856 | 56640 | 15055 | 94.5% | Drukewitz et<br>al. 2018 | SRX2501518 |
| *Laphystia*<br>*limatula* | Asilidae | 60777554 | 30019 | 20511 | 25% | Dikow et al.<br>2017 | SRX2213169 |
| *Machimus*<br>*arthriticus* | Asilidae | 83421201 | 69849 | 16916 | 90% | Drukewitz et<br>al. 2018 | SRX2501520 |
| *Mydas clavatus* | Mydi-<br>dae | 90390602 | 54643 | 26033 | 63% | Dikow et al.<br>2017 | SRX2212335 |
| *Philonicus*<br>*albiceps* | Asilidae | 107425636 | 46977 | 27750 | 90% | Dikow et al.<br>2017 | SRX2222448 |
| *Proctacanthus*<br>*coquilletti* | Asilidae | 21978654 | 56925 | 36770 | 79% | Dikow et al.<br>2017 | SRX2213473 |
| *Scleropogon*<br>*duncani* | Asilidae | 111276014 | 50672 | 25148 | 69% | Dikow et al.<br>2017 | SRX2213475 |
| *Tolmerus*<br>*atricapillus* | Asilidae | 108444670 | 43915 | 23324 | 90% | Dikow et al.<br>2017 | SRX2213152 |

### Site-specific Signatures of Selection in Asilidae Venom Gene Families

The longest isoforms of 107,641 complete coding sequences were provided as input for FUSTr (v1.0). This identified 60,727 gene families, of which 348 contained [?] 15 sequences (the minimum number allowing enough power for subsequent analyses). Of those 348 families, 77 contained at least one amino acid site under strong positive selection. FUSTr clustered the 308 venom-annotated genes and their homologs into 151 families, about 30% of which are singletons (i.e., gene families consisting of only one sequence). Sequences annotated as one of the 14 described asilidins by DEA/WEA were clustered into 34 families. Of those with more than two sequences, only asilidin5, asilidin7, asilidin12, and asilidin13 were recovered as individual, monophyletic families. Asilidin1, asilidin2, and asilidin10 were split the most - with 5, 10, and 4 separate families respectively. Asilidin14 was not included in analyses because Transdecoder (v3.0.1) could not find a complete ORF.

Twenty three DEA/WEA-annotated venom families were large enough to be analyzed for evidence of selection. Eight of these families (~34%) were found to be under positive selection. These contain homologs to asilidin2 and asilidin11 (both described from *D. genitalis*), dehydrogenase, deaminase, several peptidases, and alpha amylase. Nine additional putative venom gene families identified by toxify were also found to be under positive selection (see below for more details). These putative venom families under positive selection are shown in Figure 1. Venom dehydrogenase (family_2127) and peptidase M13 (family_696) are disregarded because they did not contain signal peptides, a necessary prerequisite for secreted proteins. Only two toxify-predicted families are shown.

**Figure 1:**
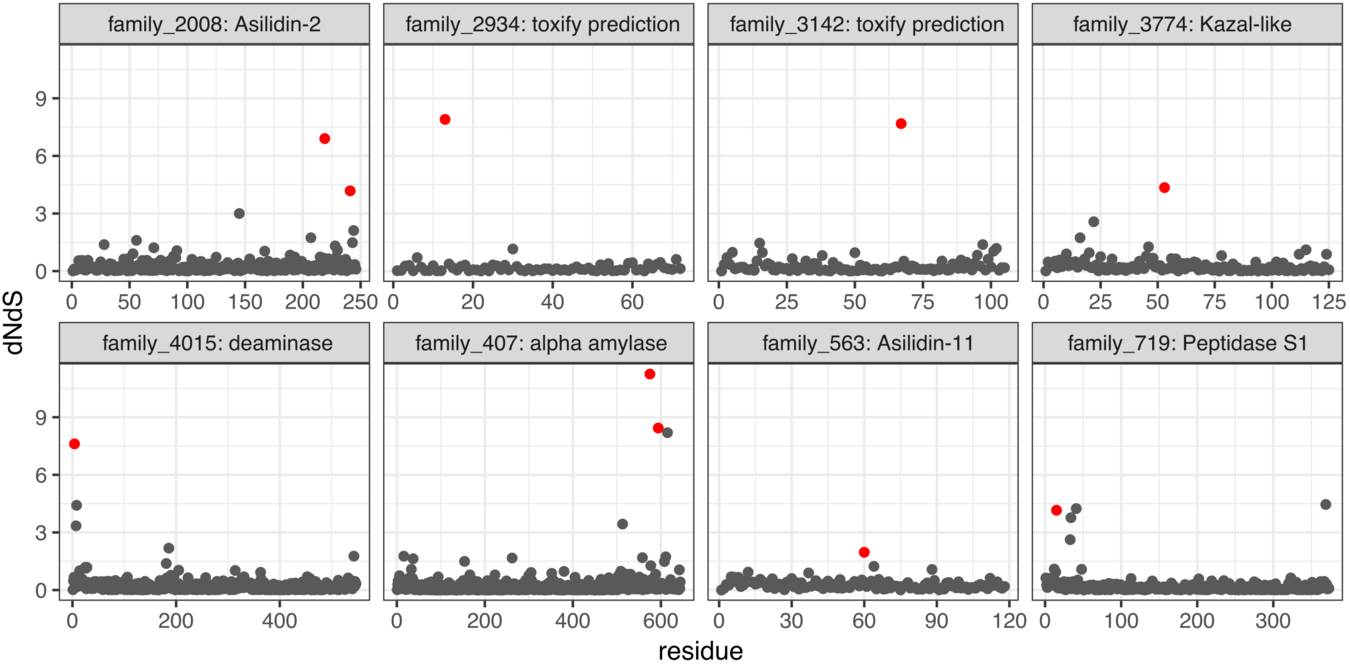
Site-specific dN/dS ratios per amino acid position for eight putative venom gene families identified via DEA/WEA and toxify. Significant sites under positive selection are marked in red.

A total of 12,976 coding sequences of the 107,641 provided as input for FUSTr were also classified as being a venom by toxify (v0.1.78) with a probability [?] 0.90. Of the 348 gene families (with [?] 15 sequences) analyzed by FUSTr, toxify classified 33 as having at least one protein sequence with a venom probability [?] 0.90, and a total of nine of those families contained at least one amino acid site under positive selection. The nonvenomous outgroups, *Apiocera parkeri* and *Mydas clavatus*, were represented in 10 and 8 families of the 33 respectively. Furthermore, toxify classified 43 of 311 annotated proteins (22 of 151 putative gene families) from *E. rufibarbis, M. arthriticus*, and *D. genitalis* as venom proteins with a probability greater than 90% (e.g., Asilidin1, Figure 2).

**Figure 2:**
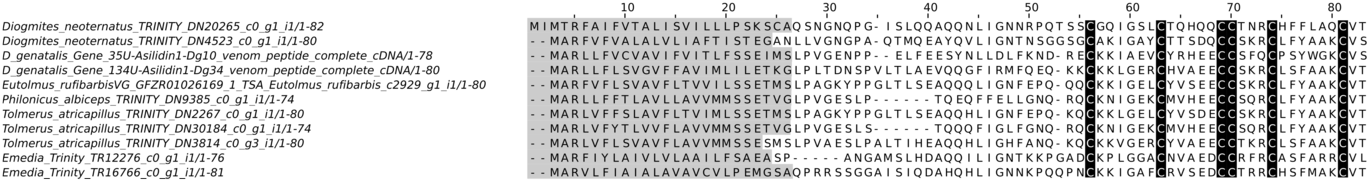
MSA of FUSTr family 2511, putative Asilidin1. Signal sequences is highlighted in grey. Cysteine residues are highlighted in black.

### Gene Family Gain/Loss in Insects

The phylogeny inferred by STAG (v1.0) is fully consistent with the topology produced by Dikow et al. (2017). The family Asilidae uniquely gained 2,509 gene families, while the sister lineage Apioceridae + Mydidae gained 27 (Figure 3). The subfamily Asilinae, from which most recent venomic studies have been conducted (e.g. [15]; [22]), gained 251 gene families. Forty-three venom-annotated (DEA/WEA) gene families were recovered as present in the ancestor of Asiloidea, while forty venom-annotated gene families were gained in Asilidae. The subfamily Asilinae gained four venom-annotated gene families. No evidence for whole-genome duplications was recovered in any lineages comprising the Asiloidea.

**Figure 3:**
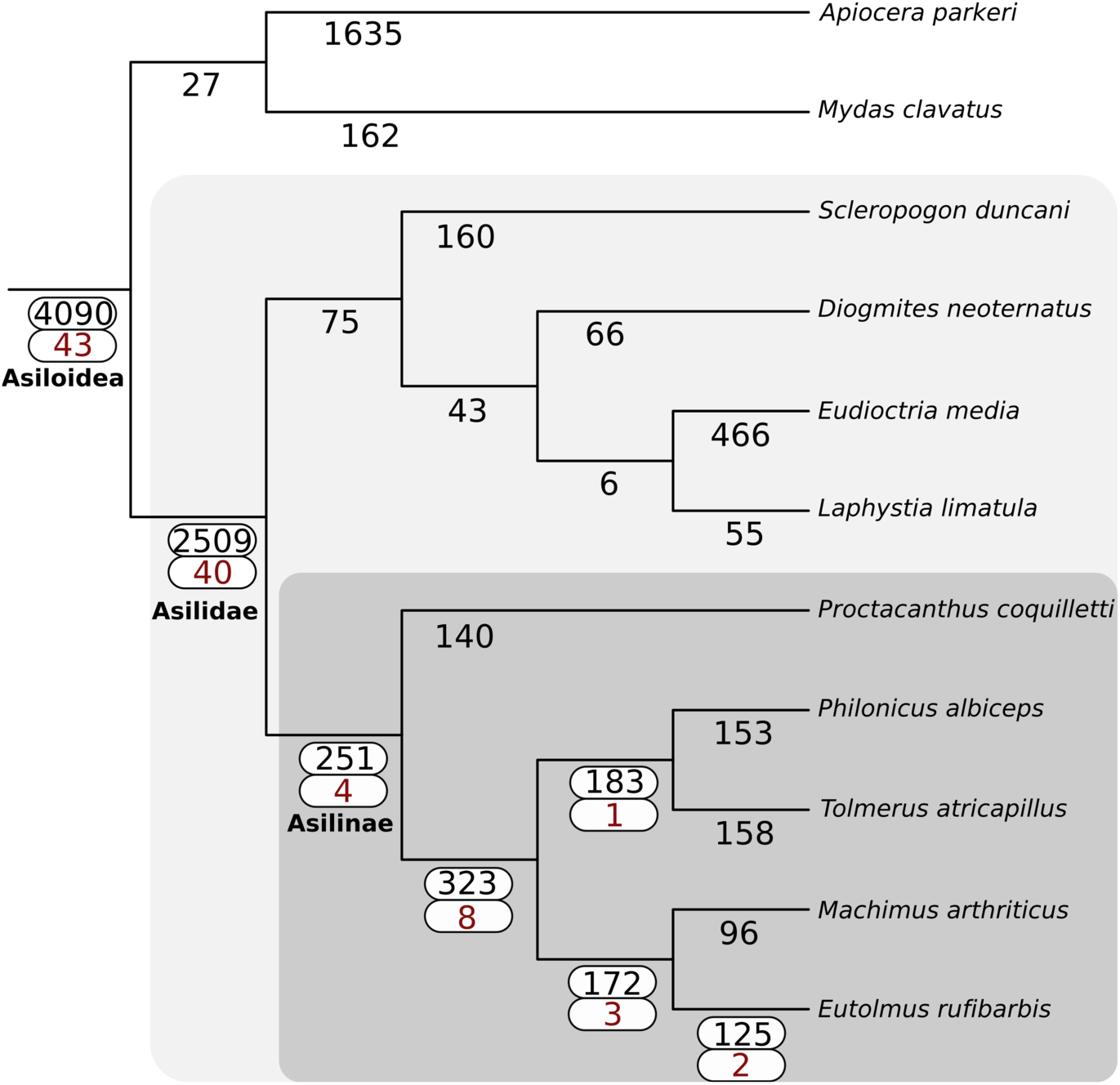
Gain of gene families across Asiloidea using SiLiX clustering and the principle of dollo parsimony as implemented in DOLLOP. The total number of gene families gained at each node are shown in black. For nodes with one or more DEA/WEA venom-annotated gene families gained (red), both values are highlighted with a white box.

## Discussion

In this study, we used FUSTr to generate putative gene families derived from thoracic and whole-body transcriptomes and to identify which of these families had evidence of positive selection. We also used toxify to assign venom probabilities to these genes, and this is the first study of putative venom genes from across the family Asilidae. Previous studies have focused on only one or two species (e.g. [15]; [22]; [23]), but in this study we included representatives from five subfamilies (including Asilinae), though this is still only a small fraction of the 14 currently recognized. We also, for the first time, examined venom gene family diversity in a broader phylogenetic context by including two representatives of the nonvenomous sister lineage to robber flies, Apioceridae + Mydidae.

### Delimitation of asilidin protein families

FUSTr split most asilidins (all except 5, 7, 12, and 13) into multiple families, suggesting that these may not be natural groups as defined by DEA and WEA. Both authors relied primarily on BLAST homology to delimit their respective asilidin families. However, it should be noted that the clustering algorithms implemented by FUSTr (i.e., SiLiX v1.2.11) have strict requirements for family assignments to reduce domain chaining, so families tend to have fewer sequences and tend to be more numerous. While we did not include data from Drukewitz et al. [23], we noticed that those authors named new asilidin families (11-15), although asilidins 11-14 had already been described by Walker et al. [22]. This confused nomenclatural situation will need to be addressed in the future.

### toxify as a complement to traditional venomics methods

This study shows that toxify can be a useful tool for identifying putative venom genes in understudied taxa for which existing genomic and/or proteomic resources are lacking. However, it should be considered as a complement to traditional venomics methods (i.e., identify candidate genes for further study), not a replacement. In Asiloidea, toxify appears to have both a high false negative and a high false positive rate. For example, only 14% of putative asilid venoms identified by previous authors [15]; [22] were predicted to be venoms by toxify. Conversely, of the gene families gained by the nonvenomous outgroups, *A. parkeri* and *M. clavatus*, 442 and 44 were predicted by toxify to contain venom sequences, respectively (Table S1).

These results may be explained by the fact that the training data for toxify consisted largely of venoms from spiders, cone snails, and snakes, and this bias may limit its effectiveness outside of those groups. As more venoms from a wider array of animals become well-characterized, toxify will be retrained, and its performance will likely improve.

### Origin of venom in Asilidae

Many putative venom gene families were already present in the ancestor of Asiloidea (41.5%), but a similar number were uniquely gained in Asilidae (39.6%). In contrast, many fewer (1.0-7.9%) venom gene families were gained in the various lineages of Asilidae studied here. This suggests that robber flies may use a suite of venoms that are fairly evolutionarily conserved across the family, supplemented by a small number of toxins unique to particular lineages. Drukewitz et al. found that the vast majority of asilid venom gene families were gained prior to the evolution of Asilidae [23], in conflict with our results. This may be due to their expanded outgroup sampling, our expanded ingroup sampling, or inherent differences in clustering methods (Orthofinder (version unreported) vs. SiLiX (v1.2.11)).

### Patterns of selection in venom proteins

The family Asilidae is roughly equivalent in age to the toxicoferan lizards included in Sunagar & Moran, which had two out of six toxin families (33%) with sites under significant positive selection [31]. For comparison, eight of twenty-three (34%) of DEA/WES-annotated robber fly venom gene families showed evidence of positive selection. This indicates that venom gene sequence evolution in asilids, like that of other ancient venomous lineages, is dominated by purifying rather than positive selection.

### Future Directions

Future venom studies in Asilidae should focus on transcriptomic and proteomic sequencing from phylogenetically disparate species, as well as toxicity assays of individual venom proteins. In addition, whole genome sequencing of a phylogenetically diverse array of robber flies and their relatives will be necessary to properly explore the genomic processes involved [6]; [23]. Only with all these resources available will we be able to gain a truly comprehensive understanding of asilid venom diversity and its evolution.

## Materials and Methods

All analyses use default parameters unless otherwise noted, and results are available via DOI: 10.6084/m9.figshare.13130171.

### Taxon sampling

Eight high-quality thoracic asiloid transcriptomes and two asilid venom gland transcriptomes were downloaded from the NCBI Short Read Archive (see Table 1 for accession numbers). To this we added an additional whole body transcriptome from *Eudioctria media* (Asilidae: Dioctriinae) (NCBI SAMN13178732). This species was chosen because it represents a distinct lineage that is only distantly related to previously studied taxa [37]. Summary statistics for the retrieved transcriptomes can be found in Table 1. Transcriptomic data for *Dolopus genitalis* [22] was not available. Instead, 123 putative venom proteins sequences from this species were included in analyses of site specific signatures of selection. Two other available robber-fly transcriptomes representing important lineages (*Nicocles dives* and *Lasiopogon cinctus*) were not included because of low read counts and low BUSCO complete scores [33]. Venom gland transcriptomes of *Dasypogon diadema* [23] were not included because a close relative (*Diogmites neoternatus*) was already included.

### RNA isolation, sequencing, and processing

Whole body RNA was extracted from one adult male *E. media* using TRIzol^®^ (Life Technologies, Carlsbad, CA) followed by purification using the Qiagen RNeasy kit (Qiagen, Valencia, CA). The extraction was sent to the Genomic Services Lab at HudsonAlpha (Huntsville, AL) for library preparation and sequencing (100 bp; paired-end) on an Illumina HiSeq 2500.

Raw FASTQ files from all transcriptomes were provided as input for trimgalore v0.3.7 [38] to trim low quality reads and remove adapters. Trimmed reads were then assembled in Trinity v2.0.6 [39]. The program BUSCO v1.1b1 [40] was used to assess the transcriptome completeness of the assembly with the included Arthropoda dataset. Protein coding sequences were then extracted from the assembled transcripts using Transdecoder v3.0.1 [41].

### Site-specific Signatures of Selection in Asilidae Venom Gene Families

In order to characterize the molecular evolution of the robber-fly venom genes described by DEA/WEA, transcriptome assemblies from nine Asilidae and two outgroups within Asiloidea (*Mydas clavatus* and *Apiocera parkeri*), as well as 123 putative venoms from WEA, were provided as input for FUSTr v1.0 [35] to detect gene families undergoing pervasive positive selection. FUSTr accomplished this by performing tests on codon alignments with the reconstructed phylogenies of gene families. Sequences were clustered into gene families using SiLiX (v1.2.11) [42]. To maintain minimum statistical power, only gene families with 15 or more sequences were analyzed for signatures of selection.

### Venom protein family assignment

Because nearly all of the described venom protein sequences from DEA/WEA were included in this analysis, we were able to classify families as probable “venoms” if they contained homologs to the previously described asilidin venom proteins. Additionally, we described novel putative venom gene families with toxify v0.1.78 [36], which uses a deep learning approach to infer whether a given protein sequence is a toxin. Putative venom proteins were only further characterized if the presence of a signal peptide was detected using SIGNALP v5.0 [43].

### Gene family loss/gain

In order to quantify the loss and gain of gene families in Asilidae in a phylogenetic framework, the gene family trees from FUSTr were provided as input to STAG v1.0 [44]. STAG takes gene trees from any multi-copy gene family that has all sequences from all species present and estimates divergence between each species pair from the closest estimated orthologous gene pairs. Through a consensus approach STAG is then able to infer the species tree in a manner that has been demonstrated to be more accurate than multi-species coalescent-based approaches such as ASTRAL or concatenated alignment. STAG is ideal for systems that have a history of duplication events that render one-to-one ortholog-only approaches unfeasible. To test whether asilid lineages have undergone substantial gains in gene families relative to other insects, we reconstructed ancestral gain and loss events for each FUSTr family for all branches in the inferred insect phylogeny using DOLLOP v3.69 from the PHYLIP package, [45] and supplementary python scripts implemented through FUSTr (https://github.com.tijeco/FUSTr).

To investigate the origins of novel gene families and expansion of paralogs, the EvoPipes.net package DupPipe was employed on the authors’ server and results were delivered 14 December 2018. The dataset of Li *et. al* [46] was combined with the *E. media* transcriptome and the CDS sequences from the *Protocanthus coquilletti* genome [47].

## Supplementary Materials

All supplementary files are found at https://figshare.com/projects/Molecular_Evolution_of_Venom_Proteins_in_Asilidae_(Insecta:_Diptera)/71897

Table S1: CSV file with columns for sequence name, taxon, FUSTr family identity, Drukewitz et al./Walker et al. annotation, whether positive selection was detected in the family, whether the family was inferred to be novel to Asilidae, toxify prediction score, and to which phylogenetic node the taxon belongs. DOI: 10.6084/m9.figshare.10324526

FUSTr cds: complete coding sequences output by FUSTr. DOI:10.6084/m9.figshare.10324565

## Acknowledgements

We thank Dr. Scott “Woody” Fitzgerald for assistance collecting the specimen of *Eudioctria media* used in this study. We also would like to thank Dr. David Maddison for providing lab space and reagents used in the RNA extraction of *Eudioctria media*. Additionally, thanks to Dr. Michael S. Barber and Zheng Li for analyzing our data on a local installation of the EvoPipes.net DupPipes package.

Funding was provided by East Carolina University Department of Biology via M.S.B’s startup.

## Author Contributions

Conceptualization: T.J.C., C.M.C. and M.S.B.; Methodology: T.J.C., C.M.C. and M.S.B.; Software: T.J.C.; Validation: C.M.C.; Formal Analysis: T.J.C. and C.M.C.; Investigation: C.M.C.; Resources: C.M.C and M.S.B.; Data Curation: T.J.C. and C.M.C.; Writing - Original Draft Preparation: C.M.C. and T.J.C.; Writing - Review & Editing: C.M.C., T.J.C., and M.S.B.; Visualization: T.J.C. and C.M.C; Supervision: M.S.B.; Project Administration: C.M.C. and M.S.B.; Funding Acquisition: M.S.B.

## Conflicts of Interest

The authors declare no conflict of interest.

